# Dopamine D1-like receptor blockade and stimulation decreases operant responding for nicotine and food in male and female rats

**DOI:** 10.1101/2022.01.24.477590

**Authors:** Ranjithkumar Chellian, Azin Behnood-Rod, Ryann Wilson, Karen Lin, Grace Wing-Yan King, Marcella Ruppert-Gomez, Alexandria Nicole Teter, Adriaan W. Bruijnzeel

## Abstract

Dopamine has been implicated in smoking, but there remains a need for a better understanding of the effects of dopamine D1-like receptor agonists on nicotine intake and the role of sex in the effects of dopaminergic drugs on nicotine intake. This work studied the effects of D1-like receptor stimulation and blockade on operant responding for nicotine and food and locomotor activity in male and female rats. The effects of the D1-like receptor antagonist SCH 23390 (0.003, 0.01, 0.03 mg/kg) and the D1-like receptor agonist A77636 (0.1, 0.3, 1 mg/kg) on responding for nicotine and food, and locomotor activity were investigated. The effects of SCH 23390 were investigated 15 min and 24 h after treatment, and the effects of the long-acting drug A77636 were investigated 15 min, 24 h, and 48 h later. Operant responding for nicotine and food was decreased 15 min, but not 24 h, after treatment with SCH 23390. Operant responding for nicotine was decreased 15 min, 24 h, and 48 h after treatment with A77636, and food responding was decreased 15 min and 24 h later. Locomotor activity was decreased 15 min, but not 24 h, after treatment with SCH 23390. A77636 only decreased locomotor activity 48 h after treatment. There were no sex differences in the effects of SCH 23390 or A77636. In conclusions, the D1-like receptor antagonist SCH 23390 reduces nicotine intake and causes sedation in rats. Stimulation of D1-like receptors with A77636 decreases nicotine intake at time points that the drug is not sedative.

## INTRODUCTION

The use of tobacco products has severe adverse effects on human health. People who use tobacco products are at high risk for various types of cancer and heart disease (Huxley and Woodward 2011; Sasco et al. 2004). Tobacco products and nicotine, which is the main psychoactive compound, induce mild euphoria and improves cognition (Benowitz 2009; Rezvani and Levin 2001). Nicotine addiction is characterized by the compulsive use of tobacco products and negative affective withdrawal signs upon cessation of nicotine use (American Psychiatric Association 2013; Bruijnzeel 2012). Despite a gradual decline in tobacco use, smoking remains a significant worldwide health burden (WHO 2018). In the US, there are about 40 million people who use tobacco products, and worldwide there are 1.3 billion tobacco users (CDC 2014; WHO 2021). Although the use of tobacco cigarettes is on the decline, there has been a substantial increase in the use of electronic cigarettes (e-cigarettes). E-cigarette use among high schoolers doubled from 2017 to 2019 (Miech et al. 2019). In the US, about 36 percent of 10th graders (12-13 years of age) and 40% of 12th graders (17-18 years of age) have used e-cigarettes. Furthermore, the COVID-19 pandemic has led to an increase in the use of e-cigarettes (Clendennen et al. 2021).

Dopamine plays a critical role in reward learning, motivated behavior, and positive reinforcing effects of drugs of abuse (Heymann et al. 2020; Pierce and Kumaresan 2006). Studies with humans and animals show that dopamine plays a role in the acute rewarding properties of nicotine (Benowitz 2010; Bruijnzeel 2012). Dopamine mediates its effect in the brain via D1-like subfamily (D1 and D5 receptors) and D2-like subfamily (D2, D3, and D4 receptors) receptors (Bourne 2001). Genetic studies point to a role for D1-like and D2-like receptors in smoking (Comings et al. 1996; Huang et al. 2008). Studies with rats indicate that D1-like and D2-like receptor activation contributes to the rewarding effects of nicotine (Corrigall and Coen 1991; Hall et al. 2015; Ivanova and Greenshaw 1997; O’Neill et al. 1991). Furthermore, exposure to nicotine induces adaptations in D1-like receptor function, which contribute to the anhedonia associated with nicotine withdrawal (Bruijnzeel and Markou 2004). Dopamine D1-like receptor levels are decreased in smokers, and their levels increase upon smoking cessation (Dagher et al. 2001; Yasuno et al. 2007). Systemic administration of nicotine induces dopamine release in the striatum (Nisell et al. 1994), and striatal dopamine release has widespread effects throughout the brain and plays a critical role in reward-related behaviors (Li and Jasanoff 2020).

Human studies indicate that there is a sex differences in smoking behavior. Nicotine might be more reinforcing in men than women, and smoking cues might play a greater role in smoking in women than men (Perkins et al. 2001; Perkins et al. 2002). A positron emission tomography (PET) study also showed that smoking differently affects the dopamine system of male and female smokers (Cosgrove et al. 2014). The dopaminergic response to smoking is greater in the ventral striatum of males and in the dorsal putamen of females. Some, but not all, animal studies suggest that there are sex differences in the reinforcing properties of nicotine. Female rats acquire nicotine intake faster, have a higher level of nicotine intake when the response requirements are low and are more motivated to self-administer nicotine under a progressive ratio schedule (Chellian et al. 2020a; Donny et al. 2000; Lynch 2009). In contrast, there is no evidence for sex differences in the effects of noncontingent nicotine administration or nicotine self-administration on reward thresholds in the intracranial self-stimulation procedure (Chellian et al. 2021a; Xue et al. 2018).

Despite that sex differences in the rewarding properties of nicotine have been explored, it is unknown if dopamine agonists and antagonists affect nicotine intake differently in males and females. Furthermore, it is not known if a selective D1-like receptor agonist affects operant responding for nicotine in rats. In the present studies, we investigated the role of dopamine signaling in the reinforcing properties of nicotine in male and female rats. The rats were treated with the D1-like receptor antagonist SCH 23390 or the D1-like receptor agonist A77636. We evaluated the acute and delayed effects of these drugs on nicotine intake. It was also determined if SCH 23390 and A77636 affect operant responding for food and locomotor activity.

## MATERIALS AND METHODS

### Animals

Adult male and female Wistar rats (males 200–250 g, females 175–225 g, 8-9 weeks of age) were purchased from Charles River (Raleigh, NC). Rats were housed with a rat of the same sex in a climate-controlled vivarium on a reversed 12 h light-dark cycle (light off at 7 AM). Food and water were available ad libitum in the home cage except for when the rats were allowed to respond for nicotine or food when they were fed 90-95 percent of their ad lib home cage intake. The experimental protocols were approved by the University of Florida Institutional Animal Care and Use Committee.

### Drugs

(-)-Nicotine hydrogen tartrate (Sigma-Aldrich), SCH 23390 hydrochloride (Tocris bioscience), and A77636 hydrochloride (Tocris bioscience) were dissolved in sterile saline (0.9 % sodium chloride). SCH 23390 and A77636 were administered subcutaneously (SC) in a volume of 1 ml/kg body weight. Nicotine was dissolved in sterile saline, and the rats self-administered 0.03 or 0.06 mg/kg/inf of nicotine in a volume of 0.1 ml/inf. Nicotine doses are expressed as base, and SCH 23390 and A77636 doses are expressed as salt.

### Experimental design

In the first experiment, the effects of SCH 23390 (Expt. 1A; males n=7, females n=9) and A77636 (Expt. 1B; males n=9, females n=9) on nicotine self-administration was investigated. In the second experiment, the effects of SCH 23390 (Expt. 2A; males n=8, females n=8) and A77636 (Expt. 2B; males n=8, females n=8) on operant responding for food was investigated, and in the third experiment, the effects of SCH 23390 (Expt. 3A; males n=8, females n=8) and A77636 (Expt. 3B; males n=8, females n=8) on locomotor activity was investigated. The treatment schedule was the same in experiments 1-3. Both SCH 23390 and A77636 were administered (SC) 15 min before nicotine self-administration, food responding, or the small open field test. It was also determined if SCH 23390 affected nicotine and food intake and locomotor activity 24 h after treatment and if A77636 affected these parameters 24 and 48 h after treatment. SCH 23390 (0, 0.003, 0.01, and 0.03 mg/kg), and A77636 (0, 0.1, and 0.3 mg/kg) were administered according to a Latin square design. The highest dose of A77636, 1mg/kg, was not included in the Latin square design and was administered at last. There was at least 48 h between injections with SCH 23390 and 72 h between injections with A77636.

### Experiment 1: effects of SCH 23390 and A77636 on nicotine self-administration

The rats were trained to respond for food pellets (45 mg, F0299, Bio-Serv, Frenchtown, NJ) in operant chambers (Med Associates, St. Albans, VT) under a fixed-ratio 1, time-out 10s (FR1-TO10s) schedule. After the food training sessions were completed, the rats were implanted with a catheter in the jugular vein. The rats were allowed to self-administer nicotine for nine daily 1-h sessions. The rats self-administered 0.03 mg/kg/infusion (inf) of nicotine under a FR1-TO10s schedule on days 1-3. The rats self-administered 0.06 mg/kg/inf of nicotine under a FR1-TO60s schedule on days 4-9. The effects of SCH 23390 (0, 0.003, 0.01, and 0.03 mg/kg, SC), and A77636 (0, 0.1, 0.3, and 1 mg/kg, SC) on nicotine intake were investigated in daily 1 h sessions (0.06 mg/kg/inf). The rats were fed 90-95 percent of their baseline food intake after the nicotine self-administration sessions.

### Experiment 2 and 3: effects of SCH 23390 and A77636 on food responding and motor activity

Adult male and female rats were trained to respond for food pellets (45 mg, F0299, Bio-Serv, Frenchtown, NJ) in operant chambers (Med Associates, St. Albans, VT) under an FR1-TO10s schedule. After the food training, the effects of SCH 23390 (0, 0.003, 0.01, and 0.03 mg/kg, SC) and A77636 (0, 0.1, 0.3, and 1 mg/kg, SC) on the operant responding for food pellets was studied in daily 20-min sessions under an FR1-TO10s schedule (Expt. 2). The rats were fed 90-95 percent of their baseline food intake in the home cage after operant responding for food. After the food study, the rats were fed ad-lib, and the effects SCH 23390 and A77636 on locomotor activity were investigated (Expt. 3). The rats were habituated to the small open field on three consecutive days (20-min sessions) and then the effects of SCH 23390 (0, 0.003, 0.01, and 0.03 mg/kg, SC) and A77636 (0, 0.1, 0.3, and 1 mg/kg, SC) on locomotor activity were investigated in 20-min sessions.

### Food training

Food training was conducted in the operant chambers. Responding on the right lever (RL, active lever) resulted in the delivery of a food pellet. Responding on the left lever (LL, inactive lever) was recorded but did not have scheduled consequences. The food training sessions were conducted for 10 days. Instrumental training started under an FRl-TOls reinforcement schedule for 5 days (30 min session per day). After the fifth food training session, the rats singly housed and remained singly housed for the rest of the study. On day 6, the time-out period was increased to 10 s. The rats were allowed to respond for food pellets under the FR1-TO10s schedule (20 min sessions) for 5 days. Both levers were retracted during the 10 s time-out period. During the food training period, the rats were fed 90-95 percent of their baseline food intake in the home cage.

### Intravenous catheter implantation and operant responding for nicotine

The catheters were implanted as described before (Chellian et al. 2021a; Chellian et al. 2021b). The rats were anesthetized with an isoflurane-oxygen vapor mixture (1-3%) and prepared with a catheter in the right jugular vein. The catheters consisted of polyurethane tubing (length 10 cm, inner diameter 0.64 mm, outer diameter 1.0 mm, model 3Fr, Instech Laboratories, Plymouth Meeting, PA). The right jugular vein was isolated, and the catheter was inserted to a depth of 3 cm for males and 2.5 cm for females. The tubing was then tunneled subcutaneously and connected to a vascular access button (Instech Laboratories, Plymouth Meeting, PA). The button was exteriorized through a small, 1-cm incision between the scapulae. After the surgery, the rats were given at least seven days to recover. The rats received daily IV infusions of the antibiotic Gentamycin (4 mg/kg, Sigma-Aldrich, St. Louis, MO) for seven days. A sterile heparin solution (0.1 ml, 50 U/ml) was flushed into the catheter before and after administering the antibiotic or nicotine self-administration. Then 0.05 ml of a sterile heparin/glycerol lock solution (500 U/ml) was infused into the catheter. The animals received carprofen (5 mg/kg, SC) daily for 48 hours after the surgery. The rats were allowed to self-administer nicotine for nine daily 1 h sessions. During the first three sessions, the rats self-administered 0.03 mg/kg/inf of nicotine under an FR1-TO10s schedule. For the following six sessions the rats self-administered 0.06 mg/kg/inf of nicotine under an FRl-TO6Os schedule. The 0.06 mg/kg/inf dose of nicotine is a relatively high dose and leads to high levels of nicotine intake (Chaudhri et al. 2005; Shoaib and Stolerman 1999).

During the first day of each dose the amount of nicotine that the rats could self-administer was limited to prevent aversive effects. The maximum number of infusion was set to 20 for the first day that the rats received the 0.03 mg/kg/inf dose and to 10 for the first day that the rats received the 0.06 mg/kg/inf dose. Active lever (right lever, RL) responding resulted in the delivery of a nicotine infusion (0.1 ml infused over a 5.6-s period). The initiation of the delivery of an infusion was paired with a cue light, which remained illuminated throughout the time-out period. Inactive lever (left lever, LL) responses were recorded but did not have scheduled consequences. Both levers were retracted during the time-out period. During the self-administration period, the rats received about 90-95 percent of their normal ad lib food intake in the home cage. The rats were fed immediately after the operant sessions.

### Small open field test

The small open field test was conducted as described before (Bruijnzeel et al. 2021; Chellian et al. 2021b; Chellian et al. 2020b). The small open field test was conducted to assess locomotor activity, rearing, and stereotypies. These motor behaviors were measured using an automated animal activity cage system (VersaMax Animal Activity Monitoring System, AccuScan Instruments, Columbus, OH, USA). Horizontal beam breaks and total distance traveled reflect locomotor activity, and vertical beam breaks reflect rearing. The distance traveled is dependent on the path of the animal in the open field and is therefore considered a better indicator of locomotor activity than horizontal beam breaks. Repeated interruptions of the same beam are a measure of stereotypies. The setup consisted of four animal activity cages made of clear acrylic (40 cm × 40 cm × 30 cm; L x W x H), with 16 equally spaced (2.5 cm) infrared beams across the length and width of the cage. The beams were located 2 cm above the cage floor (horizontal activity beams). An additional set of 16 infrared beams were located 14 cm above the cage floor (vertical activity beams). All beams were connected to a VersaMax analyzer, which sent information to a computer that displayed beam data through Windows-based software (VersaDat software). The small open field test was conducted in a dark room, and the cages were cleaned with a Nolvasan solution (chlorhexidine diacetate) between animals. Each rat was placed in the center of the small open field, and activity was measured for 20 min.

## RESULTS

### Experiment 1A: Effect of the D1-like receptor antagonist SCH 23390 on operant responding for nicotine

During the first three nicotine self-administration sessions (0.03 mg/kg/inf) nicotine intake and responding on the active lever decreased (Fig. S1A, Nicotine intake, Time F2,28 =11.943, P<0.001; Fig. S1B, RL, Time F2,28 =11.872, P<0.001). The sex of the rats did not affect nicotine intake or responding on the active lever (Nicotine intake, Sex F1,14=1.748, NS; Time x Sex 2,28=0.454, NS; RL, Sex F1,14=2.048, NS; Time x Sex 2,28=0.556, NS). Responding on the inactive lever did not change over time and was not affected by the sex of the rats (Fig. S1B, Time F2,28 =0.354, NS; Sex F1,14=0.011, NS; Time x Sex 2,28=1.449, NS). During the following six sessions (0.06 mg/kg/inf of nicotine), nicotine intake increased and was not affected by the sex of the rats (Fig. 1A, Time F5,70 =4.256, P<0.01; Sex F1,14=0.95, NS; Time x Sex F5,70=1.4, NS). Responding on the active lever (RL) increased over time and there was no effect of sex (Fig. S2A, Time F5,70 =4.435, P<0.01; Sex F1,14=1.06, NS; Time x Sex F5,70=1.427, NS). Responding on the inactive lever decreased and there was no effect of sex (Fig. S2A, Time F5,70 =2.604, P<0.05; Sex F1,14=0.156, NS; Time x Sex F5,70=1.456, NS).

**Figure 1.**
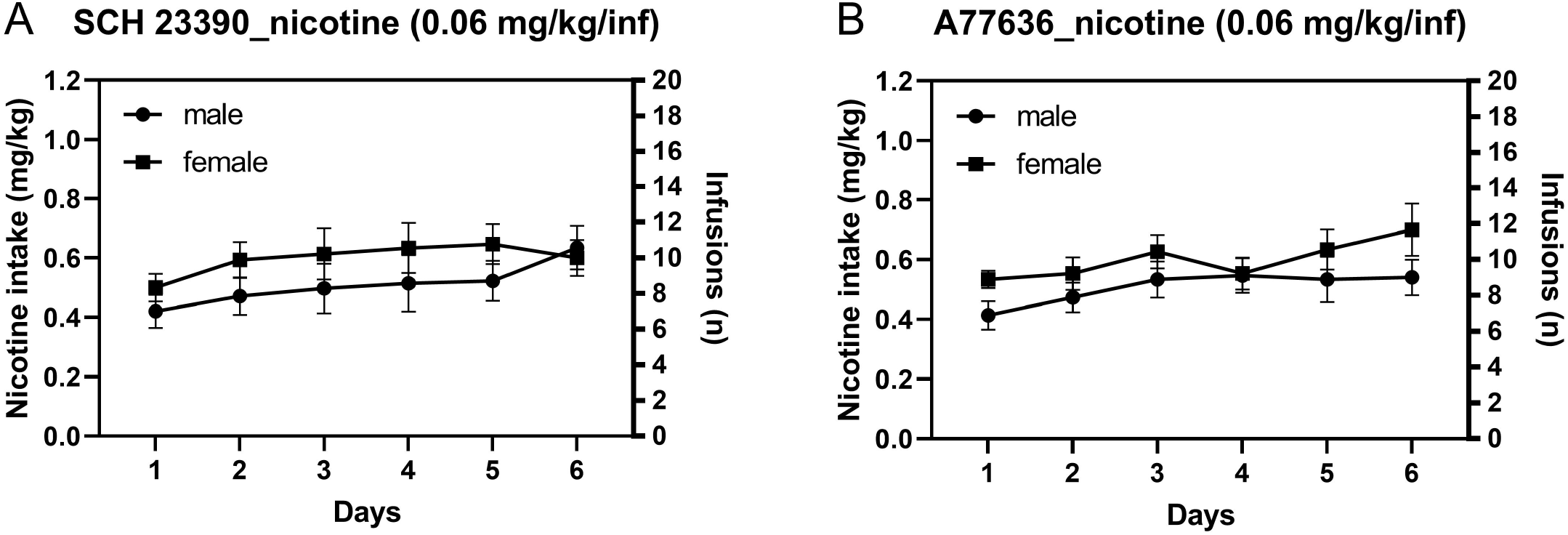
Baseline nicotine intake in male and female rats. The rats self-administered 0.06 mg/kg/inf of nicotine in 1 h sessions for six days prior to the onset of the injections with the D1 antagonist SHC 23390 (A) or the D1 agonist A77636 (B). Nicotine intake was not affected by the sex of the rats. SCH 23390 baseline group (males n=7, females, n=9), A77636 baseline group (males n=9, females n=9). Data are expressed as means ± SEM.

#### 15-min post SCH 23390 treatment

Nicotine intake was decreased 15-min after treatment with the D1-like receptor antagonist SCH 23390 and there was no effect of sex on nicotine intake (Fig. 2A, Treatment F3,42=16.932, P<0.001; Sex F1,14=3.947, NS; Treatment x Sex F3,42=1.084, NS). Responding on the active lever (RL) was decreased after treatment with SCH 23390 and the females had more active lever presses than the males (Fig. S3A, Treatment F3,42=19.312, P<0.001; Sex F1,14=5.676, P<0.05; Treatment x Sex F3,42=0.694, NS). Responding on the inactive lever (LL) was decreased after treatment with SCH 23390 and there was no effect of sex (Fig. S3C, Treatment F3,42=4.352, P<0.01; Sex F1,14=0.554, NS; Treatment x Sex F3,42=0.614, NS).

**Figure 2.**
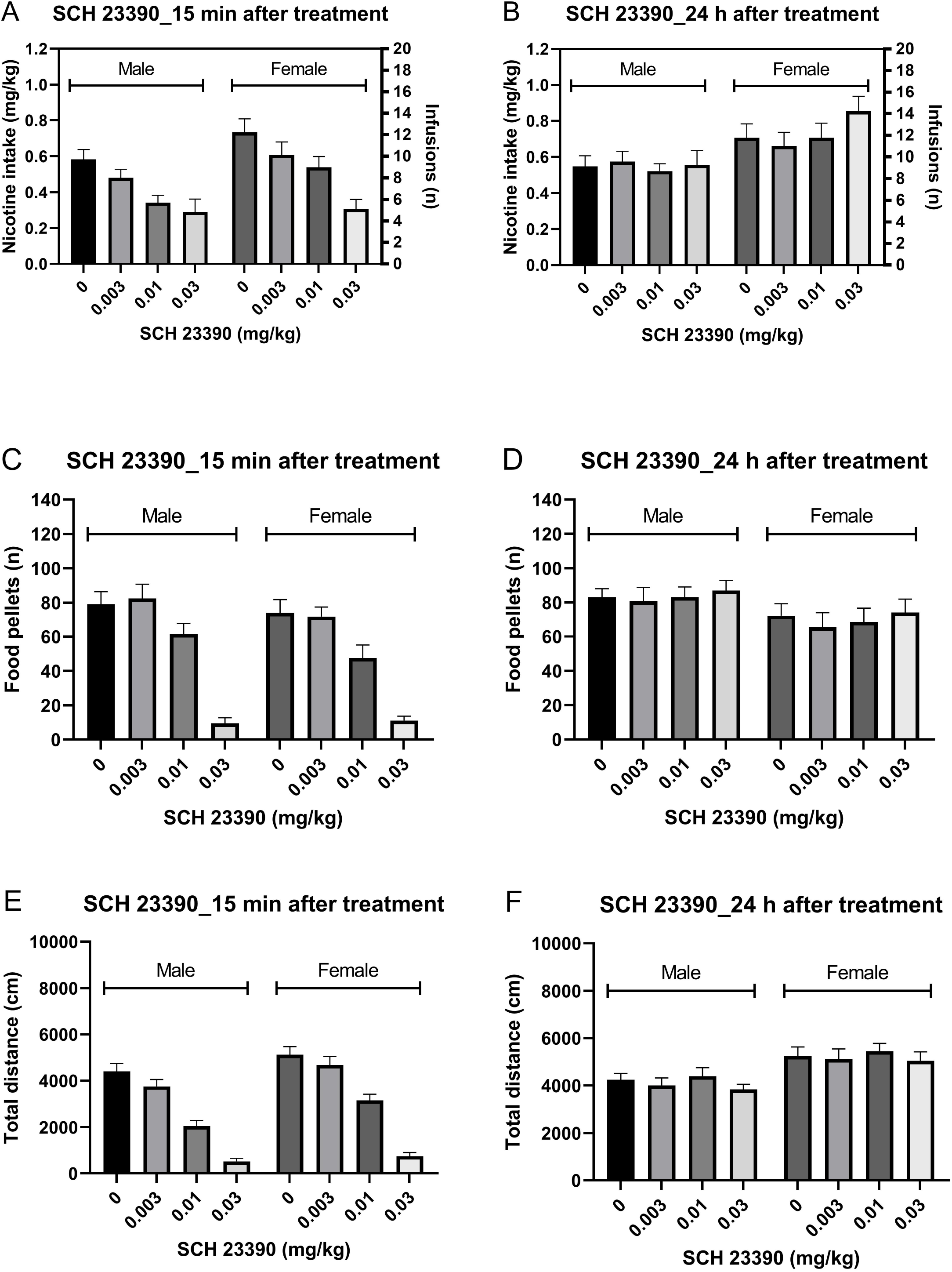
Effects of the D1-like receptor antagonist SCH 23390 on nicotine and food intake and locomotor activity in male and female rats. The effects of SCH 23390 on nicotine (A, 15 min; B, 24 h) and food intake (C, 15 min; D, 24 h) and locomotor activity (E, 15 min; F, 24 h) was investigated. Treatment with SCH 23390 decreased nicotine and food intake and locomotor activity 15 min, but not 24 h, later. Nicotine self-administration (males n=7, females, n=9), operant responding for food and small open field test (males n=8, females, n=8). Data are expressed as means ± SEM.

#### 24-h post SCH 23390 treatment

Treatment with SCH 23390 did not affect nicotine intake 24 h later and there was no effect of sex (Fig. 2B, Treatment F3,42=2.145, NS; Sex F1,14=3.908, NS; Treatment x Sex F3,42=2.206, NS). Responding on the active lever (RL) was not affected 24-h after treatment and there was no effect of sex (Fig. S3B, Treatment F3,42=1.614, NS; Sex F1,14=3.685, NS; Treatment x Sex F3,42=1.614, NS). Responding on the inactive lever (LL) was not affected after treatment and there was no sex effect (Fig. S3D, Treatment F3,42=1.183, NS; Sex F1, 14=1.431, NS; Treatment x Sex F3,42=1.323, NS).

### Experiment IB: Effect of the D1-like receptor agonist A77636 on operant responding for nicotine

During the first three nicotine self-administration sessions (0.03 mg/kg/inf) nicotine intake and responding on the active lever decreased (Fig. S1C, Nicotine intake, Time F2,32 =10.617, P<0.001; Fig. S1B, RL, Time F2,32 =8.986, P<0.001). The sex of the rats did not affect nicotine intake or responding on the active lever (Nicotine intake: Sex F1,16=1.81, NS; Time x Sex 2,32=1.427, NS; RL: Sex F1,16=1.982, NS; Time x Sex 2,32=1.415, NS). Responding on the inactive lever did not change over time, and the females responded more on the inactive lever than the males (Fig. S1D, Time F2,32=2.524, NS; Sex F1,16=7.014, P<0.05; Time x Sex 2,32=1.661, NS). During the following six sessions (0.06 mg/kg/inf) nicotine intake increased and this was not affected by the sex of the rats (Fig. 1B, Time F5,80 =3.748, P<0.01; Sex F1,16=1.871, NS; Time x Sex F5,80=0.859, NS). Responding on the active lever (RL) also increased over time, and there was no sex effect (Fig. S2B, Time F5,80 =4.607, P<0.01; Sex F1,16=1.9, NS; Time x Sex F5,80=1.041, NS). Responding on the inactive lever increased over time (Time F5,80 =3.69, P<0.01), and the females responded more on the inactive lever than the males (Sex F1,16=6.934, P<0.05; Time x Sex F5,80=1.085, NS).

It was also determined if there was a sex difference in nicotine intake when the baseline data from the first (SCH 23390) and second experiment (A77636) were combined (prior to testing the D1-like receptor agonist and antagonist). There was a no sex difference in nicotine intake during the first three sessions when the rats self-administered 0.03 mg/kg/inf of nicotine (Fig. S4, Sex F1,32=2.934, NS; Time x Sex=F2,64=0.199, NS). There was also no sex difference in nicotine intake during the following six days when the rats had access to 0.06 mg/kg/inf of nicotine (Fig. S4, Sex F1,32=2.854, NS; Time x Sex=F5,160=0.294, NS). Nicotine intake decreased during the first three sessions (0.03 mg/kg/inf; Time F2,64=17.221, P<0.001) and increased during the following six sessions (0.06 mg/kg/inf; Time F5,160=7.513, P<0.001). When days 1-9 were analyzed, there was a trend towards a higher level of nicotine intake in the females (Time F8,256=8.183, P<0.001; Sex F1,32=3.567, P=0.068; Time x Sex=F8,256=0.238, NS).

#### 15-min post A77636 treatment

Nicotine intake was decreased 15-min after treatment with A77636 and there was no effect of sex on nicotine intake (Fig. 3A; Treatment F3,48=3.631, P<0.05; Sex F1,16=1.799, NS; Treatment x Sex F3,48=1.444, NS). Responding on the active lever was also decreased after treatment with A77636 and there was no effect of sex (Fig. S5A, Treatment F3,48=3.244, P<0.05; Sex F1,16=1.787, NS; Treatment x Sex F3,48=1.4, NS). Treatment with A77636 or sex did not affect responding on the inactive lever (Fig. S5D, Treatment F3,48=1.472, NS; Sex F1,16=0.361, NS; Treatment xSex F3,48=2.711, NS).

**Figure 3.**
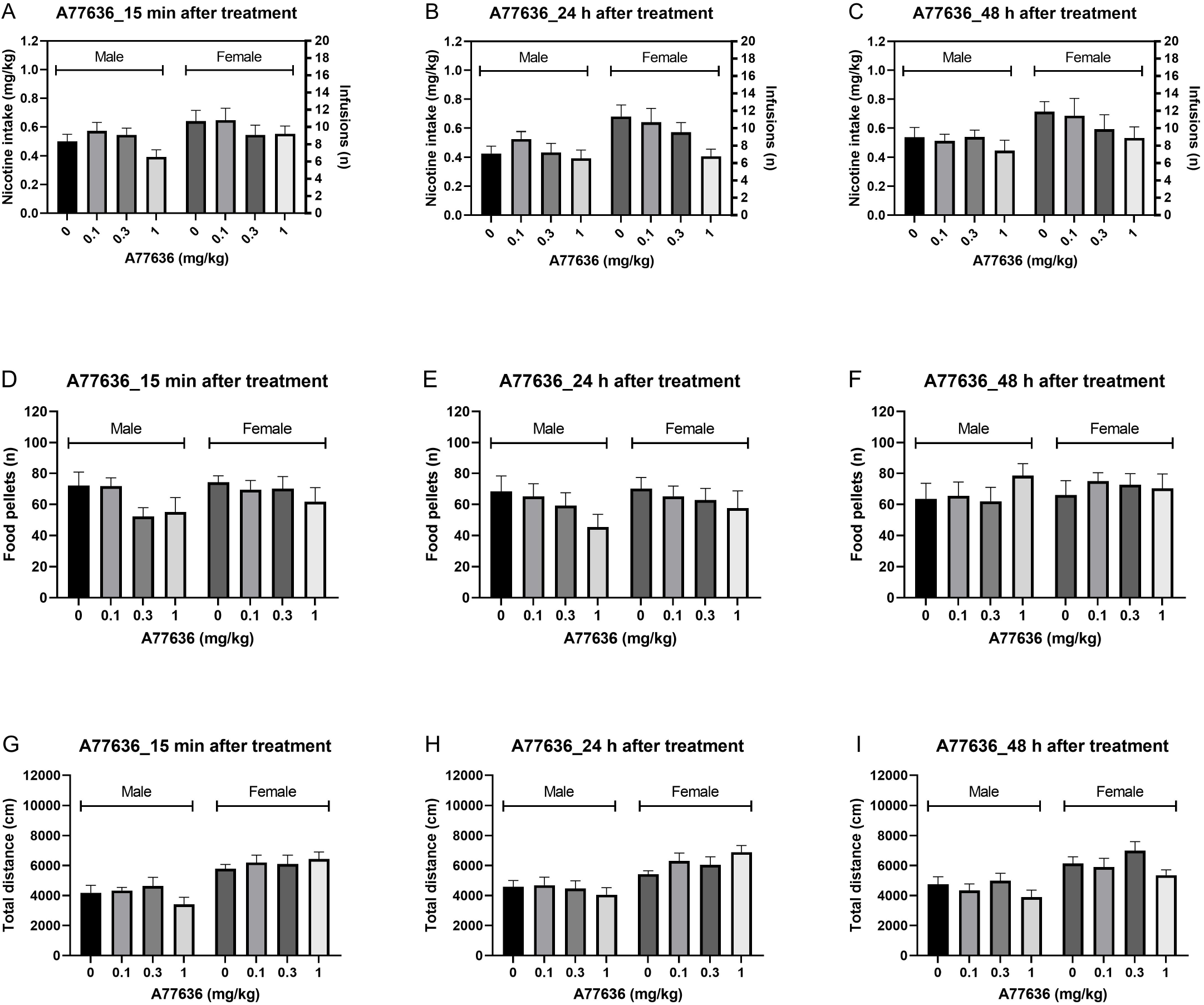
Effects of the D1-like receptor agonist A77636 on nicotine and food intake and locomotor activity in male and female rats. The effects of A77636 on nicotine (A, 15 min; B, 24 h; C, 48 h) and food intake (D, 15 min; E, 24 h; F, 48 h) and locomotor activity (G, 15 min; H, 24 h; I, 48 h) was investigated. Treatment with A77636 decreased nicotine intake 15 min, 24 h, and 48 h later. A77636 decreased food intake at the 15 min and 24 h time point and locomotor activity at the 48 h time point. Nicotine self-administration (males n=9, females, n=9), operant responding for food and small open field test (males n=8, females, n=8). Data are expressed as means ± SEM.

#### 24-h post A77636 treatment

Nicotine intake was decreased 24 h after treatment with A77636 and nicotine intake was not affected by the sex of the rats (Fig. 3B, Treatment F3,48=5.857, P<0.05; Sex F1,16=3.137, NS; Treatment x Sex F3,48=2.204, NS). Responding on the active lever was decreased after treatment with A77636 and there was no sex effect (Fig. S5B, Treatment F3,48=5.341, P<0.01; Sex F1,16=3.302, NS; Treatment x Sex F3,48=2.516, NS). Treatment with A77636 or sex did not affect responding on the inactive lever (Fig. S5E, Treatment F3,48=1.557, NS; Sex F1,16=0.758, NS; Treatment x Sex F3,48=1.018, NS).

#### 48-h post A77636 treatment

Nicotine intake was decreased 48 h after treatment with A77636 and there was no effect of sex on nicotine intake (Fig. 3C, Treatment F3,48=3.527, P<0.05; Sex F1,16=1.639, NS; Treatment x Sex F3,48=0.942, NS). Responding on the active lever was also decreased after treatment with A77636 and there was no sex effect (Fig. S5C, Treatment F3,48=2.883, P<0.05; Sex F1,16=1.855, NS; Treatment x Sex F3,48=0.476, NS). Treatment with A77636 or sex did not affect responding on the inactive lever (Fig. S5F, Treatment F3,48=0.125, NS; Sex F1,16=0.006, NS; Treatment x Sex F3,48=2.287, NS).

### Experiment 2A: Effect of the D1-like receptor antagonist SCH 23390 on operant responding for food

#### 15-min post SCH 23390 treatment

The number of food pellets received was decreased 15 min after treatment with SCH 23390 and the number of food pellets received was not affected by the sex of the rats (Fig. 2C, Treatment F3,42=68.047, p<0.001; Sex F1,14=1.23, NS; Treatment x Sex F3,42=0.812, NS). Responding on the active lever was also decreased after treatment with SCH 23390 and not affected by the sex of the rats (Fig. S6A, Treatment F3,42=68.002, P<0.001; Sex F1,14=0.821, NS; Treatment x Sex F3,42=0.887, NS). Treatment with SCH 23390 or the sex of the rats did not affect responding on the inactive lever (Fig. S6C, Treatment F3,42=2.232, NS; Sex F1,14=3.515, NS; Treatment x Sex F3,42=1.6, NS).

#### 24-h post SCH 23390 treatment

There was no effect of SCH 23390 or sex on the number of food pellets received (Fig. 2D, Treatment F3,42=0.849, NS; Treatment x Sex F3,42=0.082, NS; Sex F1,14=2.611, NS), or active lever responding (Fig. S6B, Treatment F3,42=0.866, NS; Sex F1,14=2.168, NS; Treatment x Sex F3,42=0.083, NS). Treatment with SCH 23390 did not affect inactive lever responding but the females had more inactive lever responses than the males (Fig. S6D, Treatment F3,42=1.939, NS; Sex F1,14=6.63, P<0.05; Treatment x Sex F3,42=0.083, NS).

### Experiment 2B: Effect of the D1-like receptor agonist A77636 on operant responding for food

#### 15-min post A77636 treatment

The number of food pellets received was decreased 15 min after treatment with A77636 and not affected by the sex of the rats (Fig. 3D, Treatment F3,42=3.271, P<0.05; Treatment x Sex F3,42=1.199, NS; Sex F1,14=0.655, NS). Responding on the active lever was decreased after treatment with A77636 and not affected by the sex of the rats (Fig. S7A, Treatment F3,42=3.054, P<0.05)(Sex F1, 14=0.633, NS; Treatment x Sex F3,42=1.18, NS). Treatment with A77636 or the sex of the rats did not affect responding on the inactive lever (Fig. 7D, Treatment F3,42=2.633, NS; Sex F1,14=1.151, NS; Treatment x Sex F3,42=1.049, NS).

#### 24-h post A77636 treatment

The number of food pellets received was decreased 24 h after treatment with A77636 and there was no effect of sex on the number of food pellets received (Fig. 3E, Treatment F3,42=4.701, P<0.01; Treatment x Sex F3,42=0.584, NS; Sex F1,14=0.181, NS). Responding on the active lever was also decreased after treatment with A77636 and not affected by the sex of the rats (Fig. S7B, Treatment F3,42=4.697, P<0.01; Sex F1,14=0.16, NS; Treatment x Sex F3,42=0.564, NS). Treatment with A77636 or the sex of the rats did not affect responding on the inactive lever (Fig. S7E, Treatment F3,42=1.528, NS; Sex F1,14=0.001, NS; Treatment x Sex F3,42=0.331, NS).

#### 48-h post A77636 treatment

The number of food pellets received was not affected by treatment with A77636 or the sex of the rats (Fig 3F, Treatment F3,42=1.403, NS; Treatment x Sex F3,42=1.525, NS; Sex F1,14=0.119, NS). Responding on the active lever was not affected by treatment with A77636 or sex (Fig. S7C, Treatment F3,42=1.355, NS; Sex F1,14=0.16, NS; Treatment x Sex F3,42=1.52, NS). Treatment with A77636 or the sex of the rats did not affect responding on the inactive lever (Fig. S7F, Treatment F3,42=0.424, NS; Sex F1,14=2.157, NS; Treatment x Sex F3,42=1.075, NS).

### Experiment 3A: Effect of the D1-like receptor antagonist SCH 23390 on locomotor activity

#### 15-min post SCH 23390 treatment

Treatment with SCH 23390 decreased the total distance traveled and the females traveled a greater distance than the males (Fig. 2E, Treatment F3,42 = 107.826, P<0.001; Sex F1,14=0.767, 8.418, P<0.05; Sex × Treatment F3, 42 = 1.098, NS). Treatment with SCH 23390 also decreased horizontal beam breaks and the females had the same number of horizontal beam breaks as the males (Fig. S8A, Treatment F3,42 = 117.259, P<0.001; Sex F1,14=0.767, NS; Sex × Treatment F3, 42 = 1.772, NS). SCH 23390 decreased vertical beam breaks and the females had more vertical beam breaks than the males (Fig. S8C, Treatment F3,42 = 70.615, P<0.001; Sex F1,14=18.216, P<0.001; Sex × Treatment F3,42 = 2.358, NS). SCH 23390 decreased stereotypies and the number of stereotypies was not affected by the sex of the rats (Fig. S8E, Treatment F3,42 = 64.603, P<0.001; Sex F1,14=1.165, NS; Sex × Treatment F3, 42 = 3.929, P<0.05).

#### 24-h post SCH 23390 treatment

Treatment with SCH 23390 did not affect the total distance traveled 24 h later, and the females traveled a greater distance than the males (Fig. 2F, Treatment F3,42 = 1.196, NS; Sex F1,14=10.476, P<0.01; Sex × Treatment F3,42 = 0.049, NS). Treatment with SCH 23390 and the sex of the rats did not affect horizontal beam breaks (Fig. S8B, Treatment F3,42 = 0.527, NS; Sex F1,14=0.566, NS; Sex × Treatment F3,42= 0.168, NS). Treatment with SCH 23390 did not affect vertical beam breaks and the females had more vertical beam breaks than the males (Fig. S8D, Treatment F3,42 = 1.108, NS; Sex F1,14=15.032, P<0.01; Sex × Treatment F3,42 = 0.058, NS). SCH 23390 and the sex of the rats did not affect stereotypies (Fig. S8F, Treatment F3,42 = 0.43, NS; Sex F1,14=0.924, NS; Sex × Treatment F3,42 = 0.279, NS).

### Experiment 3B: Effect of the D1-like receptor agonist A77636 on locomotor activity

#### 15-min post A77636 treatment

Treatment with A77636 did not affect the total distance traveled and the females traveled a greater distance than the males (Fig. 3G, Treatment F3,42 = 0.583, NS; Sex F1,14=35.041, P<0.001; Sex × Treatment F3, 42 = 2.106, NS). A77636 did not affect the horizontal beam breaks but the females had more horizontal beam breaks than the males (Fig. S9A, Treatment F3,42 = 2.121, NS; Sex F1,14=6.118, P<0.05; Sex × Treatment F3, 42 = 2.387, NS). A77636 did not affect vertical beam breaks but the females had more vertical beam breaks than the males (Fig. S9D, Treatment F3,42 = 3.536, P<0.05; Sex F1,14=21.205, P<0.001; Sex × Treatment F3,42 = 1.451, NS). A77636 decreased the number of stereotypies and stereotypies were not affected by the sex of the rats (Fig. S9G, Treatment F3,42 = 3.983, P<0.05; Sex F1,14=1.376, NS; Sex × Treatment F3, 42 = 2.686, NS).

#### 24-h post A77636 treatment

A77636 did not affect the total distance traveled and the females traveled a greater distance than the males (Fig. 3H, Treatment F3,42 = 0.571, NS; Sex F1,14=13.994, P<0.01; Sex × Treatment F3, 42 = 2.763, NS). A77636 and the sex of the rats did not affect horizontal beam breaks (Fig. S9B, Treatment F3,42 = 0.285, NS; Sex F1,14=2.314, NS;). There was also a drug treatment x sex interaction but the post hoc test did not reveal significant effects (Sex × Treatment F3,42 = 3.386, P<0.05). A77636 decreased vertical beam breaks and the females had more vertical beam breaks than the males ((Fig. S9E, Treatment F3,42 = 5.752, P<0.01; Sex F1,14=7.33, P<0.05; Sex × Treatment F3, 42 = 0.186, NS). A77636 did not affect stereotypies and stereotypies were not affected by the sex of the rats (Fig. S9H, Treatment F3,42 = 0.691, NS; Sex F1,14=0.155, NS;). There was also a drug treatment x sex interaction but the post hoc test did not reveal significant effects (Sex × Treatment F3, 42 = 5.382, P<0.01).

#### 48-h post A77636 treatment

A77636 decreased the total distance traveled and the females traveled a greater distance than the males (Fig. 3I, Treatment F3,42 = 4.645, P<0.01; Sex F1,14=12.376, P<0.01; Sex × Treatment F3, 42 = 0.299, NS). Treatment with A77636 decreased horizontal beam breaks and the sex of the rats did not affect the horizontal beam breaks (Fig. S9C, Treatment F3,42 = 8.955, P<0.001; Sex F1,14=1.498, NS; Sex × Treatment F3, 42 = 0.211, NS). A77636 decreased vertical beam breaks and the females had more vertical beam breaks than the males (Fig. S9F, Treatment F3,42 = 5.51, P<0.01; Sex F1,14=7.506, P<0.05; Sex × Treatment F3, 42 = 1.052, NS). A77636 did not affect stereotypies and stereotypies were not affected by the sex of the rats (Fig. S9I, Treatment F3,42 = 2.677, NS; Sex F1,14=0.002, NS; Sex × Treatment F3, 42 = 0.71, NS).

## DISCUSSION

In the present study, we investigated the effects of the dopamine D1-like receptor antagonist SCH 23390 and the dopamine D1-like receptor agonist A77636 on operant responding for nicotine and food and locomotor activity in adult male and female rats. Nicotine intake was decreased 15 min, but not 24 h, after the administration of the D1-like receptor antagonist SCH 23390. Nicotine intake was also decreased 15 min, 24 h, and 48h after the administration of the D1-like receptor agonist A77636. Operant responding for food was decreased 15 min after the administration of SCH 23390, but not 24 h later. Food intake was also decreased 15 min and 24 h, but not 48 h, after the administration of A77636. Locomotor activity was decreased 15 min after the administration of SCH 23390 but not 24 h later. Treatment with A77636 only affected locomotor activity at the 48 h time point. The females traveled a greater distance than the males in the small open field test. The sex of the rats did not affect the effects of SCH 23390 or A77636 on nicotine and food intake and locomotor activity. The present findings show that the D1-like receptor antagonist SCH 23390 induces a brief decrease in nicotine and food intake and that the D1-like receptor agonist A77636 induces a prolonged decreased in nicotine and food intake. The D1-like receptor agonist A77636 decreased nicotine and food intake at time points (15 min and 24-h post treatment) that did not affect locomotor activity.

We investigated the effects of the D1-like receptor agonist A77636 on nicotine and food intake and locomotor activity. The D1-like receptor agonist A77636 had a long-term effect on nicotine intake. A77636 decreased nicotine intake 15 min, 24 h and 48 h after treatment. We are not aware of other studies that investigated the effects of A77636 or other D1-like receptor agonists on nicotine self-administration in rats. However, studies have investigated the effects of other D1-like receptor agonists on cocaine self-administration in rats. It has been shown that the D1-like receptor agonists SKF 77434 and SKF 82958 decrease cocaine self-administration in male Wistar rats (Caine and Koob 1994). Furthermore, low doses of A77636 that do not affect locomotor activity and decrease the cocaine-induced locomotor response in male Swiss Webster mice (Chausmer and Katz 2002). These findings suggest that D1-like receptor agonists decrease the reinforcing properties of at least some addictive psychostimulants. Several studies have investigated the effects of D1-like receptor agonists on brain reward function. The dopamine D1-like receptor agonists A77636 and SKF82958 facilitate intracranial self-stimulation (ICSS), which suggests that these compounds have rewarding properties (Carr et al. 2001; Lazenka et al. 2016). In addition, dopamine D1-like receptor agonists SKF81297, SKF82958, ABT-431 produced conditioned place preference in rats (Graham et al. 2007). However, rhesus monkeys do not self-administer the D1-like receptor agonist SKF 38393 (Woolverton et al. 1984). Furthermore, the D1-like receptor agonists SKF 82958 and SKF 77434 do not maintain IV self-administration in rats with a history of cocaine intake (Caine and Koob 1994). These findings indicate that D1-like receptor agonists decrease drug intake and have some rewarding properties. However, because the D1-like receptor agonists do not maintain self-administration they might have low abuse potential.

In the present study, we investigated the effects of SCH 23390 on the self-administration of a high dose of nicotine (0.06 mg/kg/inf) in male and female rats. Treatment with the D1-like receptor antagonist led to a large decrease in nicotine intake in the male and the female rats. The present finding is in line with a previous study that investigated the effects SCH 23390 on the self-administration of a lower dose of nicotine (0.03 mg/kg/inf) in adult male Long-Evans rats (Corrigall and Coen 1991). Furthermore, it has also been shown that SCH 23390 decreases IV nicotine self-administration in adult female Sprague-Dawley rats (Willette et al. 2019). Our work shows that SCH 23390 has a similar effect on nicotine intake in male and female rats. In the present study, we also investigated if SCH 23390 affects operant responding for food and activity in the small open field. We found that SCH 23390 decreases operant responding for food and also decreases activity in the small open field in the males and the females. The highest dose of SCH 23390 decreased nicotine intake by 50 percent but also caused a large decrease in operant responding for food and locomotor activity. The observation that SCH 23390 induces a large decrease in open field activity suggests that blockade of D1-like receptors has sedative effects or impairs motor function. Indeed, it has been reported that blockade of D1-like receptors in humans and monkeys can cause sedation (Gilbert et al. 2014; Lublin et al. 1993). Overall, the present findings indicate that blockade of D1-like receptors decreases nicotine intake but also causes a large decrease in food intake and locomotor activity. The sedative effects of SCH 23390 might mitigate its effectiveness as a smoking cessation treatment.

In the present study, we also determined sex differences in nicotine intake, food intake, and open-field behavior. There was no significant sex differences in baseline nicotine intake and there was no sex difference in nicotine intake during the experiment. In our previous work with adult Wistar rats, we observed a higher level of nicotine intake in female than male rats (Chellian et al. 2021a; Chellian et al. 2021b). However, in rats that were trained to respond on the active lever for food pellets before the nicotine self-administration sessions sex differences were only observed after the acquisition phase (10 days of nicotine self-administration)(Chellian et al. 2021a). Food training leads to a high level of nicotine intake during the acquisition phase, which may mask sex differences in nicotine intake. Other studies reported that there are no sex differences in nicotine intake when nicotine intake has been established, and the rats are tested under FR schedules with low response requirements (Chaudhri et al. 2005; Donny et al. 2000; Feltenstein et al. 2012; Grebenstein et al. 2013). Female rats may self-administer more nicotine than males during the acquisition phase in studies without prior food training (Donny et al. 2000; Park et al. 2007; Wang et al. 2018). These studies suggest that females tend to self-administer more nicotine than males and these sex differences are most likely to be observed during the acquisition phase and when the response requirements are high. A recent meta-analysis that was based on 20 studies concluded that female rats self-administer more nicotine than male rats (Flores et al. 2019). This suggests that large numbers of animals are needed to detect sex difference in nicotine intake in rats. In this study we also compared sex differences in activity in the small open field test. Compared to the males, the females traveled a greater distance and displayed more rearing. This is in line with previous studies that have shown that females rats are more active in the small open field and other behavioral tests (Bruijnzeel et al. 2021; Chellian et al. 2021b; Knight et al. 2021). We did not observe sex difference in operant responding for food pellets. This is in line with previous work in which we showed that there are no sex difference in operant responding for food pellets in adult Wistar rats when the response requirements are low (Chellian et al. 2020c). Sex difference in operant responding for food are observed when the response requirement are gradually increased using a behavioral economics procedure (Chellian et al. 2020c).

It is interesting to note that treatment with the D1-like receptor agonist A77636 had a long-term effect on nicotine and food intake. This observation is in line with other studies that showed that A77636 has long-term effects. A study with 6-hydroxydopamine (6-OHDA) lesioned animals showed that treatment with the D1-like receptor agonist A68930 causes a robust rotation response (Britton et al. 1991). However, on the second treatment day, the response was decreased by 95 percent, and on the third treatment day, the response was absent (Britton et al. 1991). This indicates that treatment with A68930 dramatically decreases the sensitivity to D1-like receptor activation. A study with SK-N-MC (neuroblastoma) cells showed that pretreatment with A68930 abolished the cAMP response to subsequent treatment with this drug (Lin et al. 1996). Furthermore, pre-incubation of rat striatal membranes with A-77636 greatly (76 percent) decreased D1-like receptor binding (Lin et al. 1996). A recent study with HEK293 cells showed that A77636 dose-dependently increases cAMP levels and decreases D1-like receptor levels by almost 50 percent (Nilson et al. 2020). Importantly, clinical studies indicate that there is a positive relationship between D1-like receptor levels and the subjective effects of drugs, and preclinical studies also show a positive association between D1 receptor levels and the behavioral responses to stimulants (Caine et al. 2007; Xu et al. 2000; Zhang et al. 2000; Zhang et al. 2021). Taken together, these studies suggest that treatment with A77636 leads to prolonged D1-like receptor desensitization and may thereby decrease the reinforcing properties of nicotine and food.

In the present study, we found that the D1-like receptor antagonist SCH 23390 and the D1-like agonist A77636 decreased operant responding for food in the male and the female rats. The D1-like agonist and antagonist had the same effect on food intake in males and females. Prior studies have reported that SCH 23390 and A77636 decrease food intake in male rats. The D1-like antagonist SCH 23390 decreases operant responding for food in male Wistar and Long Evans rats (Beninger et al. 1987; Corrigall and Coen 1991). A study with male Sprague Dawley rats found that the administration of SCH 23390 into the ventral tegmental area decreases responding for food pellets under a progressive ratio schedule (Sharf et al. 2005). Thus these studies with SCH 23390 indicate that D1-like receptors in the mesolimbic dopaminergic system plays a critical role in the reinforcing properties of food. We also found that the D1-like receptor agonist A77636 decreases food intake in male and female rats. A study with male hooded rats showed that A77636 decreased food intake by reducing the meal size and duration (Cooper et al. 2006). The effect of A77636 on food intake is most likely due to the fact that this drug causes prolonged dopamine D1-like receptor desensitization (Lin et al. 1996; Nilson et al. 2020).

In conclusion, the present findings indicate that the D1-like receptor antagonist SCH 23390 induces a brief decrease in nicotine and food intake, and the D1-like receptor agonist A77636 induces a prolonged decrease in nicotine and food intake. However, SCH 23390 doses that decreased nicotine and food intake also caused sedative effects, which may hamper the use of this drug as a smoking cessation aid. The D1-like receptor agonist A77636 decreased nicotine and food intake at doses that did not induce sedative effects. The D1-like receptor agonist A77636 could potentially be used as a smoking cessation aid or appetite suppressant.

## CRediT authorship contribution statement

**R. Chellian:** Formal analysis, Investigation, Writing – Review & Editing, Visualization. **A. Behnood-Rod:** Investigation, Project administration. **R. Wilson:** Investigation, Project administration. **K. Lin:** Investigation. **G. King:** Investigation. **M. Ruppert-Gomez:** Investigation. **A. Teter:** Investigation. **A. Bruijnzeel:** Conceptualization, Formal analysis, Writing – Original Draft, Visualization, Supervision, Project administration, Funding acquisition.

