## Supplemental Materials for "Dopamine D1-like receptor blockade and stimulation decreases operant responding for nicotine and food in male and female rats"

Figure S1

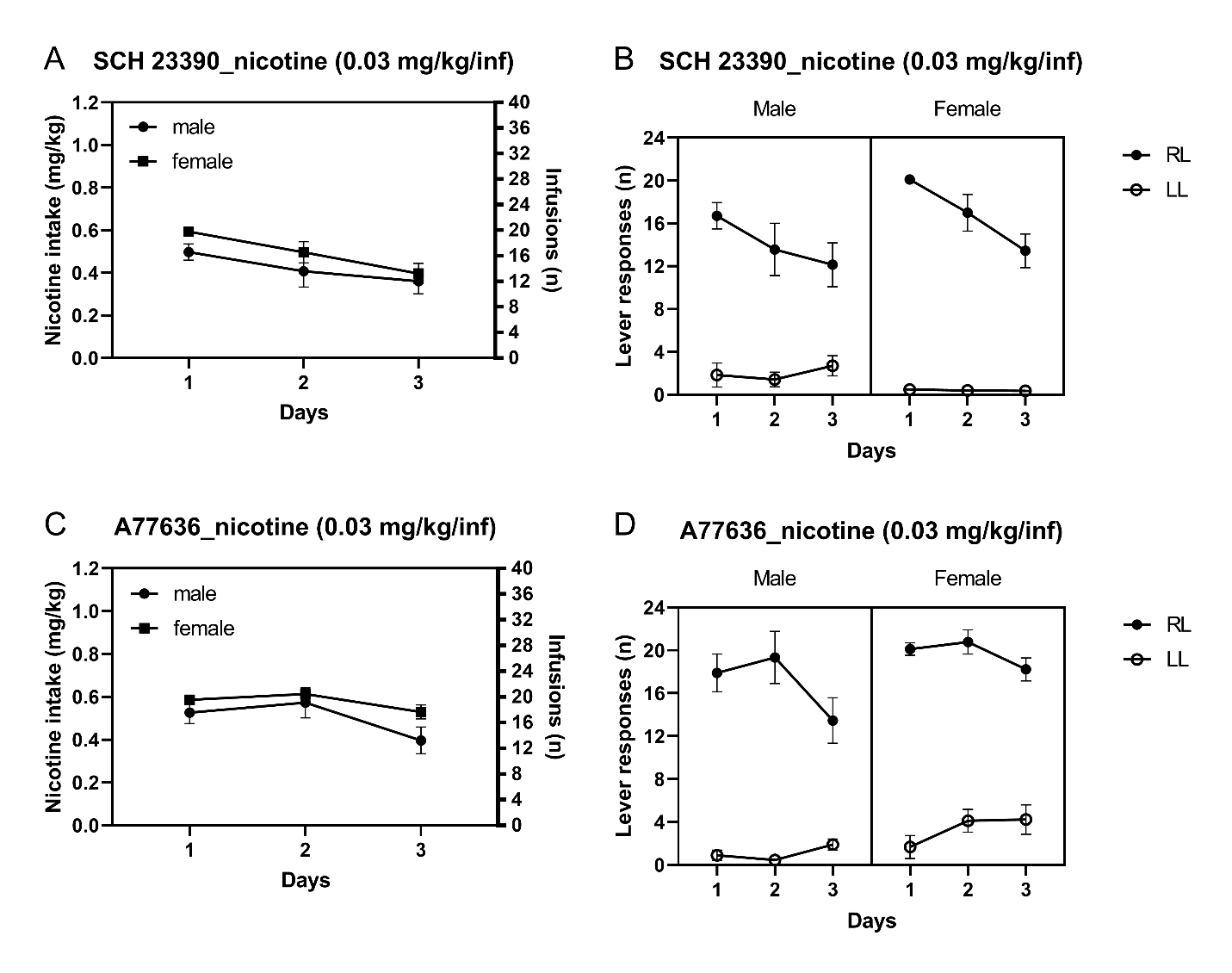

**Figure S1. Baseline nicotine intake (0.03 mg/kg/inf).** The figures depict three days of baseline nicotine intake (A) and lever responses (B) in 1 h sessions before the rats self-administered the 0.06 mg/kg/inf dose of nicotine (SCH 23390; males n=7, females, n=9). The figures also depict three days of nicotine intake (C) and lever responses (D) before the rats self-administered the 0.06 mg/kg/inf dose (A77636; males n=9, females n=9). Data are expressed as means ± SEM.

Figure S2

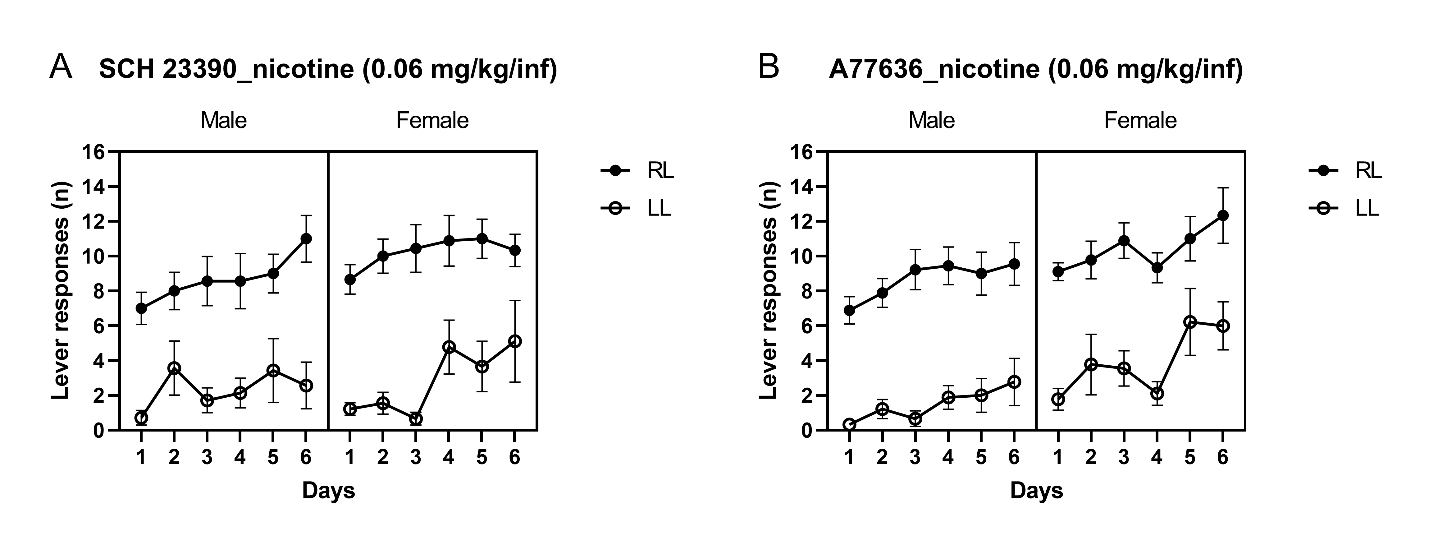

**Figure S2. Baseline nicotine intake (0.06 mg/kg/inf).** The figure depicts six days of lever responses (A, B) in 1 h sessions before treatments with SCH 23390 or A77636 started. A, males n=7, females, n=9; B, males n=9, females n=9. Data are expressed as means ± SEM.

Figure S3

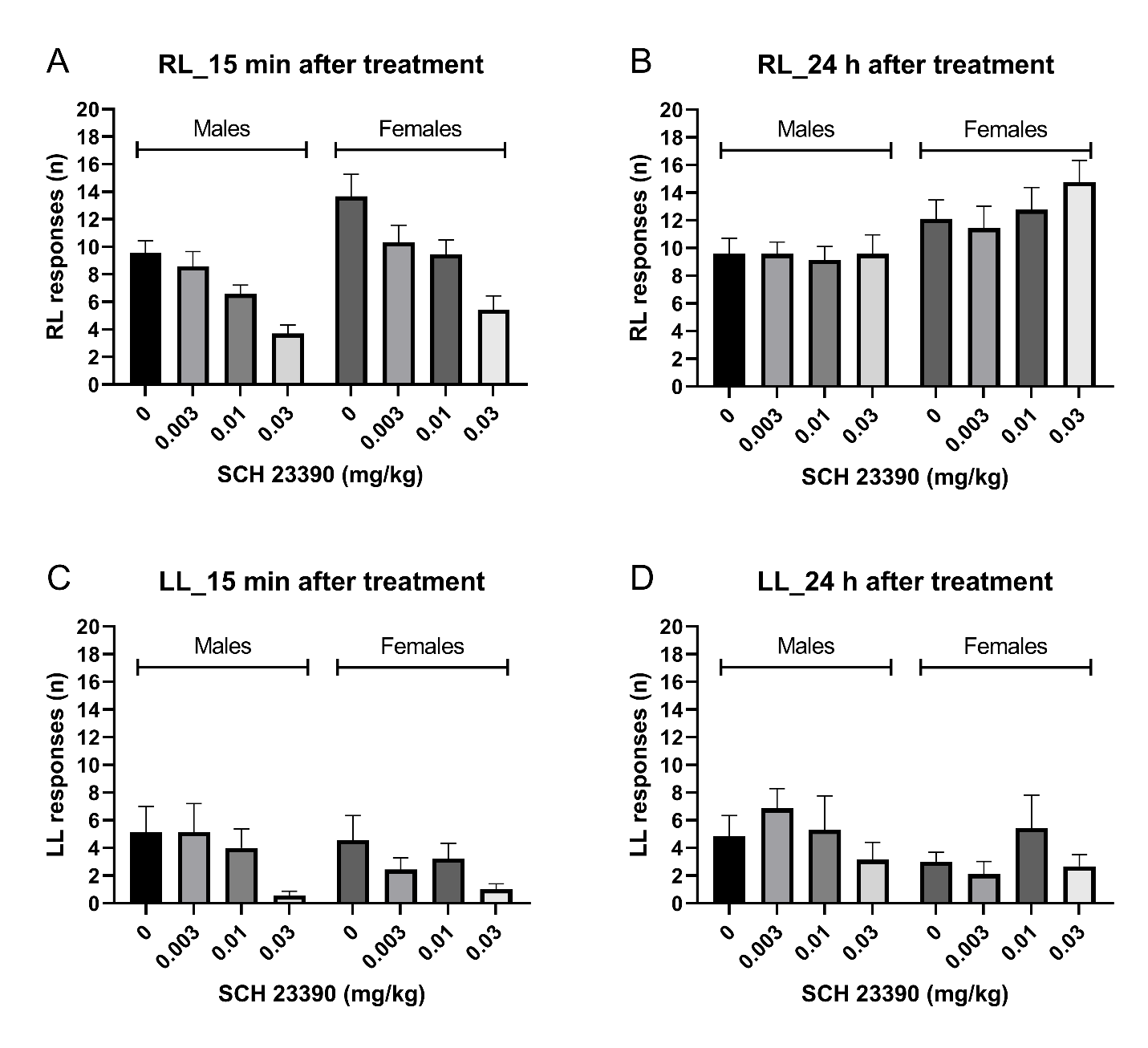

**Figure S3. Operant responding for nicotine after treatment with SCH 23390.** The figures depict responding on the active lever (A, B) and the inactive lever (C, D) 15 min and 24 h after treatment with SCH 23390. Males n=7, females n=9. Data are expressed as means ± SEM.

Figure S4

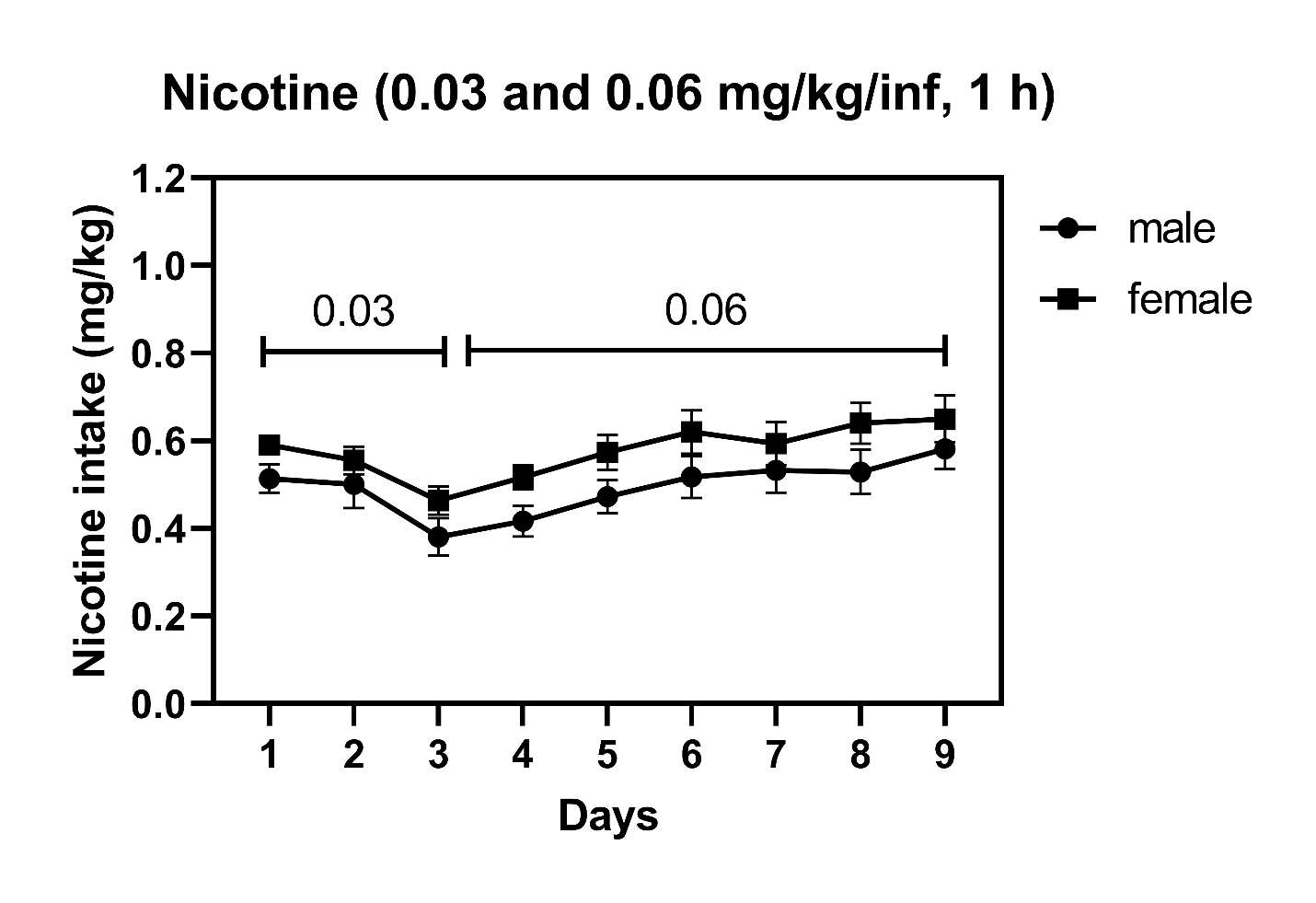

**Figure S4. Baseline nicotine intake.** The figures shows nine days of baseline nicotine intake in 1 h sessions before the onset of the drug treatments. The rats self-administered 0.03 mg/kg/inf of nicotine for three days and 0.06 mg/kg/inf of nicotine for six days. The baseline nicotine intake data from the first and second nicotine self-administration experiment were combined. Males n=16, females n=18. Data are expressed as means ± SEM.

Figure S5

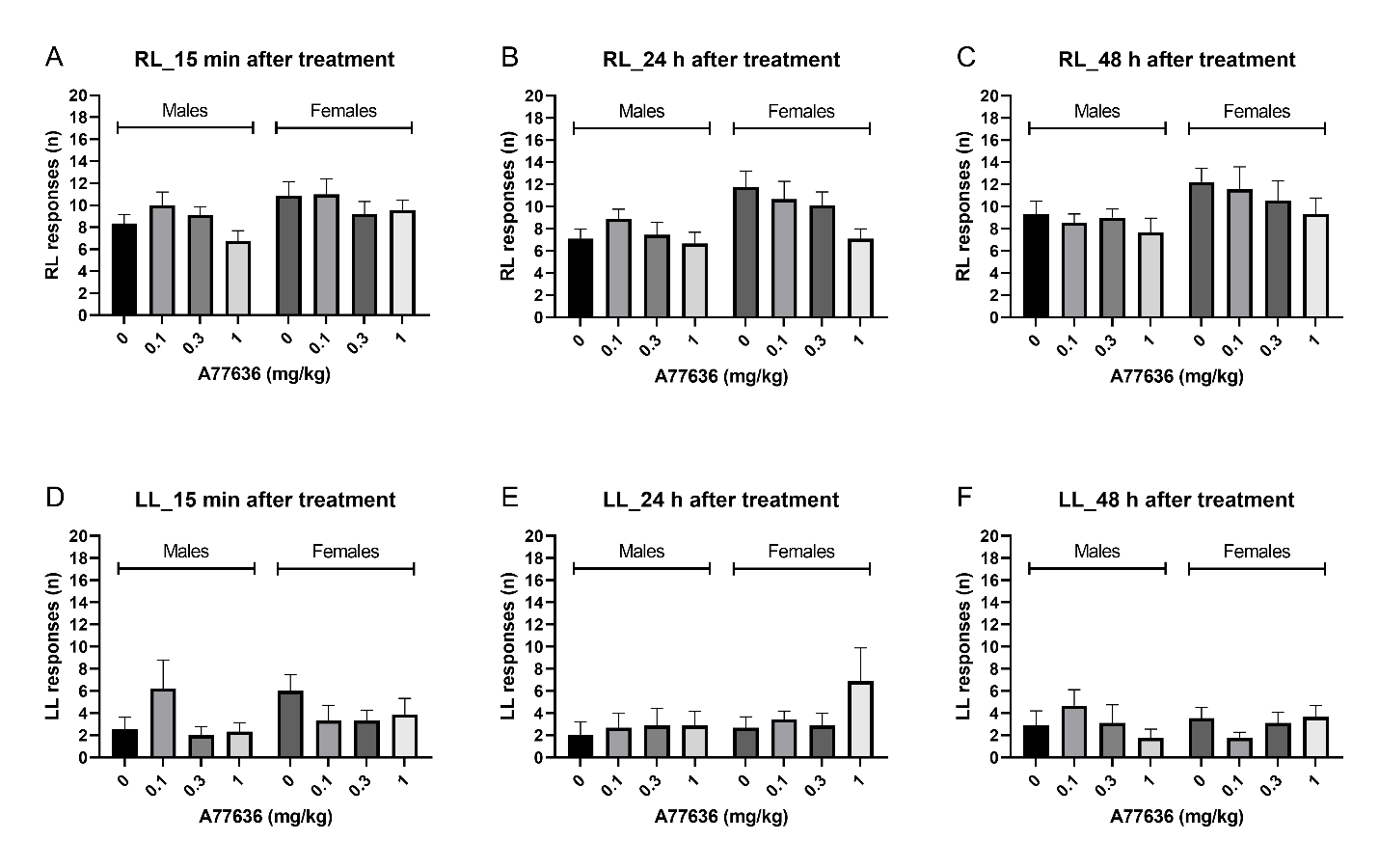

**Figure S5. Operant responding for nicotine after treatment with A77636.** The figures depict responding on the active (A, B, C) and the inactive lever (C, D, E) 15 min, 24 h, and 48 h after treatment with A77636. Males n=9, females n=9. Data are expressed as means ± SEM.

Figure S6

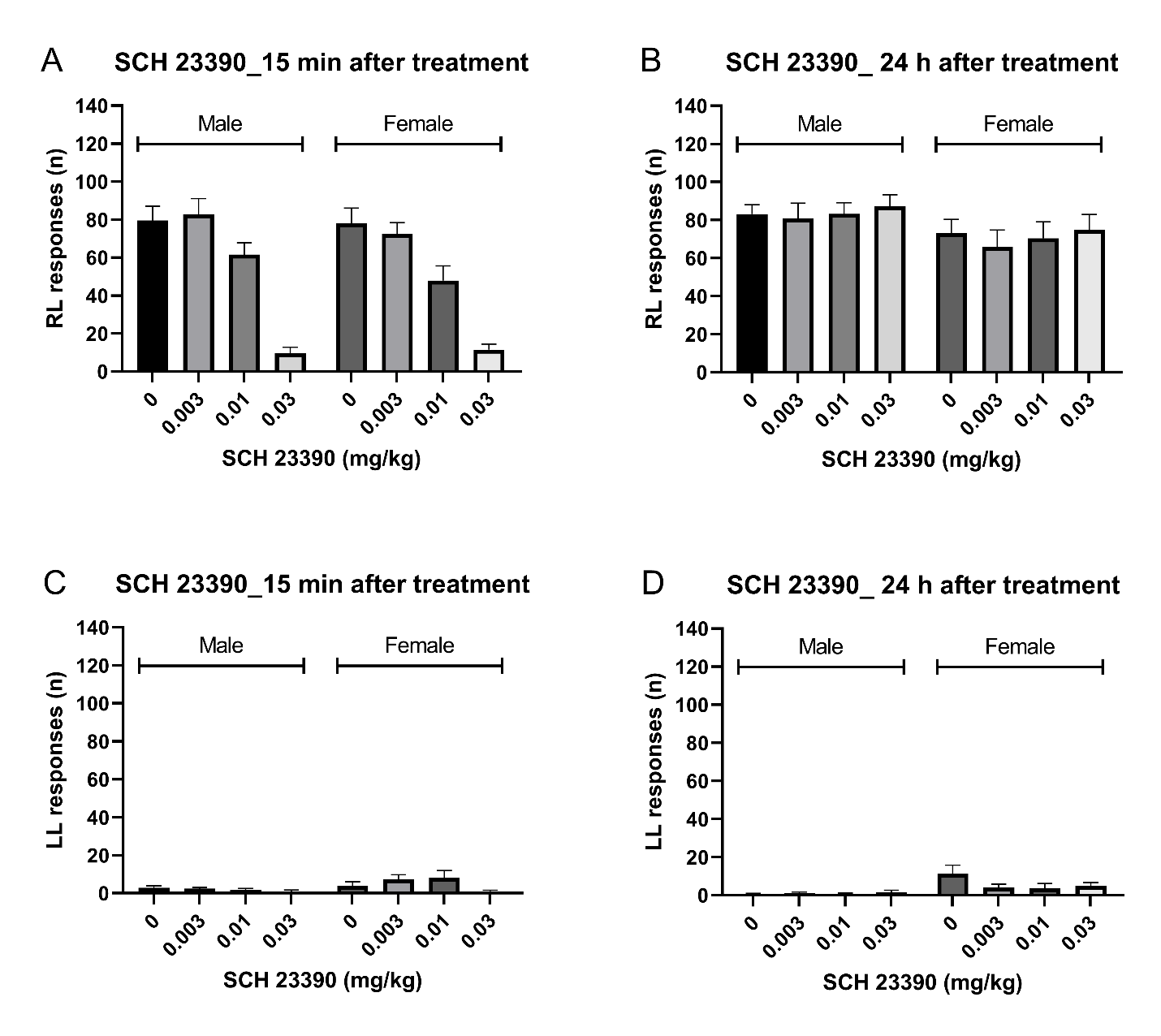

**Figure S6. Operant responding for food 15 min and 24 h after treatment with SCH 23390.** The figures depict responding on the active (A, B) and the inactive lever (C, D) 15 min and 24 h after treatment with SCH 23390. Males n=8, females n=8. Data are expressed as means ± SEM.

Figure S7

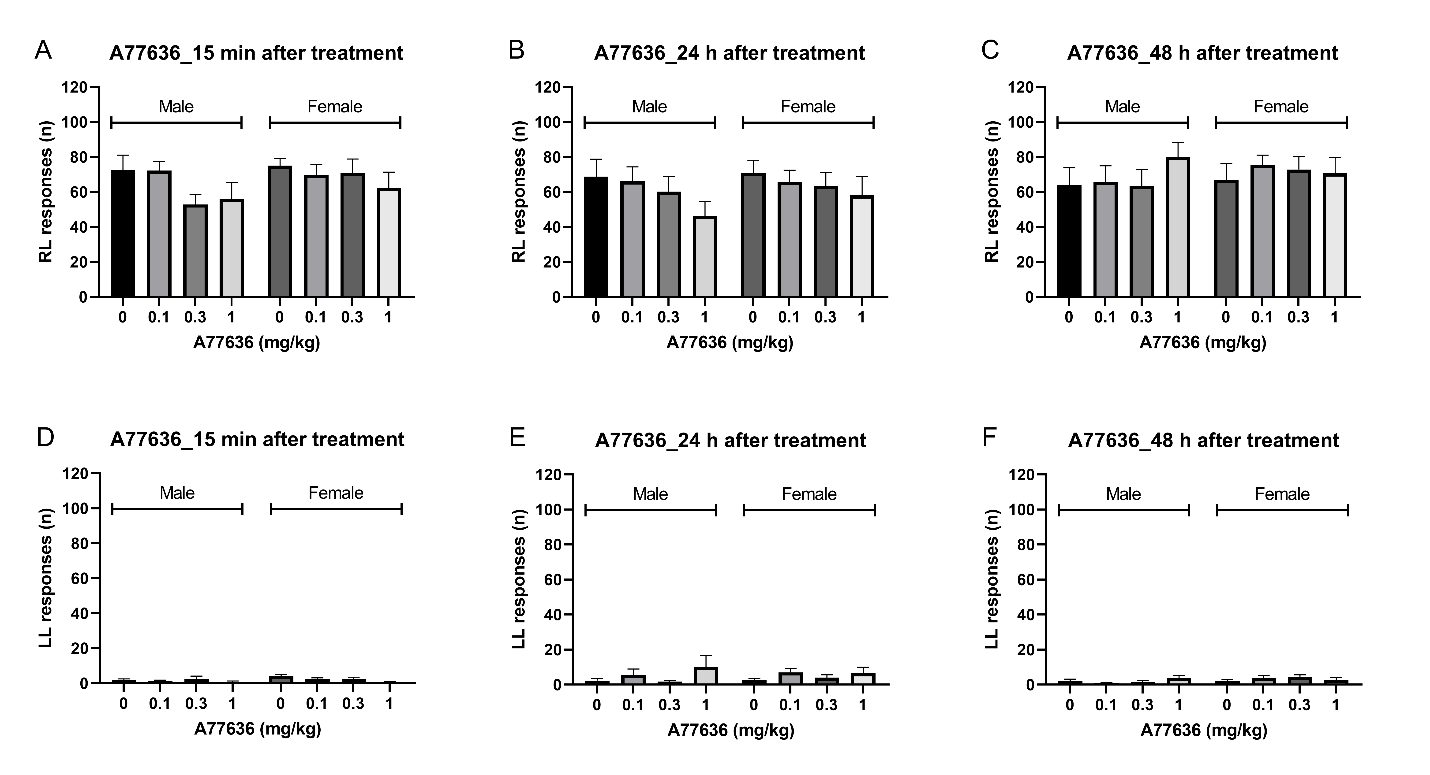

**Figure S7. Operant responding for food 15 min, 24 h, and 48 h after treatment with A77636.** The figures depict responding on the active (A, B, C) and inactive lever (D, E, F) 15 min, 24 h, and 48 h after treatment with A77636. Males n=8, females n=8. Data are expressed as means ± SEM.

Figure S8

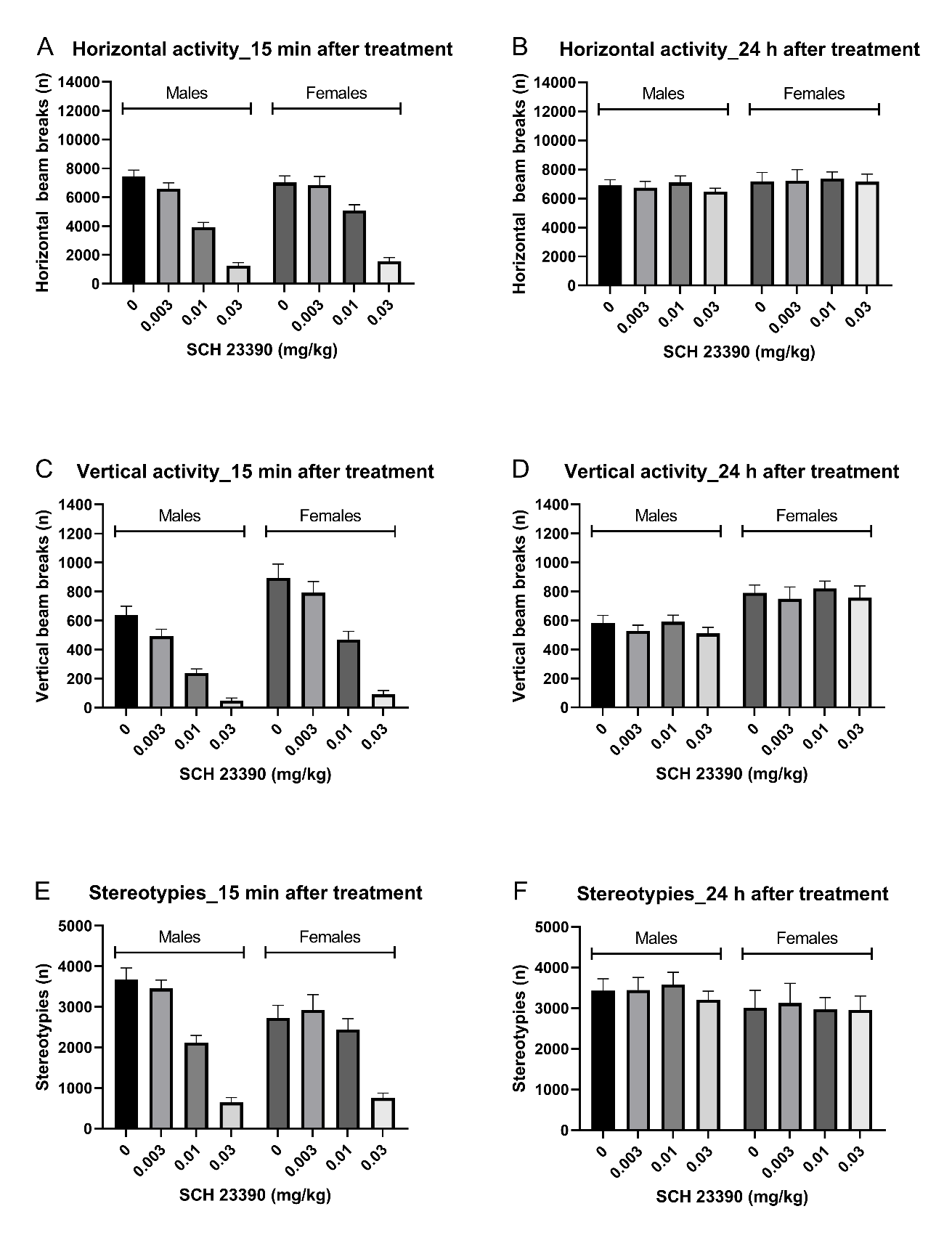

**Figure S8. Behavior in the small open field 15 min and 24 h after treatment with SCH 23390.** The figures depict horizontal beam breaks (A, B), vertical beam breaks (C, D), and stereotypies (E, F) 15 min and 24 h after treatment with SCH 23390. Males n=8, females n=8. Data are expressed as means ± SEM.

Figure S9

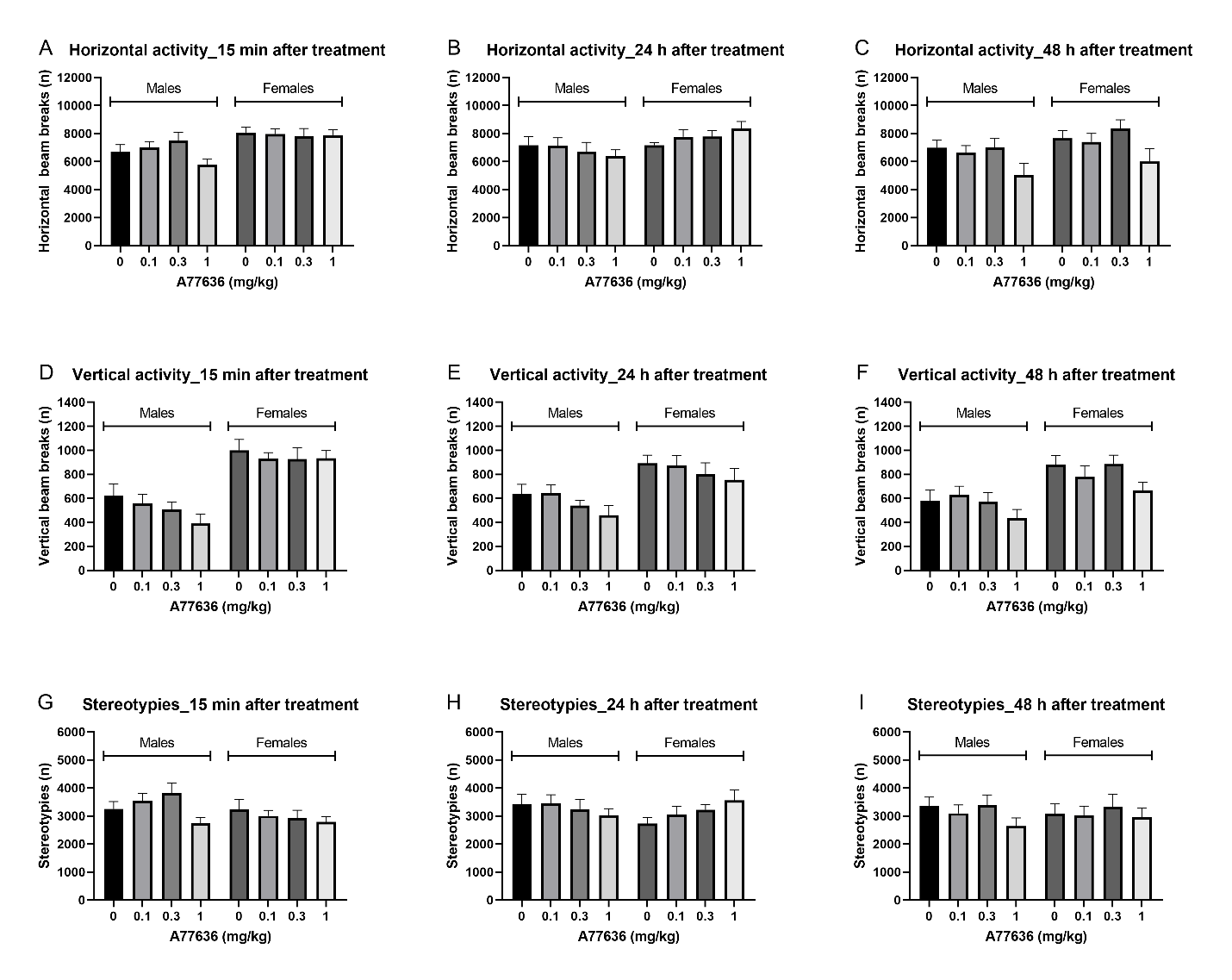

**Figure S9. Behavior in the small open field 15 min, 24 h, and 48 h after treatment with A77636.** The figures depict horizontal beam breaks (A, B, C), vertical beam breaks (D, E, F), and stereotypies (G, H, I) 15 min, 24 h, and 48 h after treatment with A77636. Males n=8, females n=8. Data are expressed as means ± SEM.
